# Label-Free Lymphocyte Subtype and State Classification by Stimulated Raman Scattering Microscopy

**DOI:** 10.64898/2026.11.16.766877

**Authors:** King Wai Chiu, Paul J. Leanza, Ziang Li, Bryce Manifold, Hao Yuan Kueh, Dan Fu

**Author notes:** **Corresponding Author** Dan Fu –.

## Abstract

Accurate immune cell subtype and functional state profiling are essential for studying immune regulation, disease progression, and therapeutic response. Established single-cell analysis techniques rely heavily on either endpoint, destructive measurements or complex labeling that can potentially affect cell functions. Stimulated Raman scattering (SRS) microscopy offers quantitative, label-free imaging of cells and tissues by providing contrast based on subcellular concentrations of intrinsic biomolecules at high spatial and temporal resolution. Here, we demonstrate that Vision Transformer based hyperspectral SRS imaging classification can accurately and non-destructively classify various lymphocyte phenotypes, including lymphocyte lineage (T and B cells), T cell subset and differentiation state (CD4^+^ and CD8^+^ T cells, effector, memory, and naïve for both), and functional states (naïve T, activated T, exhausted T) using fluorescent markers for training. By leveraging attention-based spatial-spectral feature learning, the approach captures biomolecular patterns associated with immune-cell identity and state while preserving cells for downstream or longitudinal analysis. This provides a potential platform for label-free immune profiling, live-cell analysis, and longitudinal studies of immune cell type and state transitions.

## INTRODUCTION

Cells in the body vary widely in size, shape, and subcellular features.^1^ Morphological differences, including enlarged nuclei, an increased nuclear-to-cytoplasmic ratio, and abnormal chromatin distribution, are key to identifying and differentiating cells in pathology.^2,3^ Outside pathology, morphology is also frequently used to classify cell types.^4,5^ Moreover, different functional states of the same cell type could exhibit distinct morphological features.^6–8^ Dynamic shape changes can link morphological changes to alterations in cell state and metabolism.^9–11^ However, morphological differences of many cell types are often subtle and ambiguous, such as immune cells. T and B lym-phocytes are both round, with similar cell diameter and nuclear morphology.^12^ T cells can be further categorized by function: cytotoxic T cells (CD8^+^) and helper T cells (CD4^+^). Each category can be further separated into naïve, memory, and effector cells based on their differentiation states. These cell types and states cannot be distinguished easily using morphology alone. Yet, separating these phenotypes is critical for understanding immune function and enabling biomedical applications through immune engineering. Exhausted or dysfunctional cells can reflect chronic stimulation in cancer, persistent infection, or inflammatory disease. Chimeric antigen receptor T cells (CAR-T cells) used in therapy suffer from exhaustion.^13^

The current gold-standard method for lymphocyte classification relies on immunofluorescence (IF). IF is an immunochemical technique that conjugates fluorescent dyes to primary or secondary antibodies to enable high-sensitivity detection and localization of a wide range of antigens in various cell or tissue preparations. An example application of IF is flow cytometry sorting of blood cells, where subsets of immune cells are identified and sorted based on their unique expression of surface markers.^14^ However, it is only applicable to isolated cells with established surface markers.

Cells can also be classified based on their intrinsic metabolic and biochemical signatures. For example, fluorescence lifetime microscopy (FLIM) of metabolic enzymes NADH and FAD+ reports the metabolic state of cells.^10^ It has been widely used to distinguish cancer cells from normal cells. Recently, FLIM has also been shown to classify the subtype and/or activation state of immune cells.^15–18^ Raman spectroscopy is another commonly used technique to detect the biochemical composition of cells and differentiate cell types or cell states. Subtle changes in Raman spectra of lipid and protein composition can distinguish different developmental stages of stem cells and classify immune cells.^19–22^ Spontaneous Raman in the fingerprint region was used to classify T/B cells and naïve and activated T cells.^23^ A recent report demonstrated further classification of subtypes, such as CD4 and CD8 T cells, with around 60% accuracy.^24^ Conventional T cell (Tconv) and regulatory T cells (Treg) classification has also been reported.^25^ However, Raman measurements suffer from low-throughput data acquisition. Because of time limitations in practical applications, researchers typically sample only a single point or a few points from each cell and measure only a limited number of cells, which increases spectral variability. Moreover, Raman-only measurements lack morphological information.

Stimulated Raman scattering (SRS) microscopy is an advanced Raman technique that offers high-speed chemical imaging, thus providing both morphological and chemical information.^26,27^ SRS microscopy targets the sample’s endogenous vibrational chemical differences, potentially offering a significant advantage in cell classification without the concerns of modulating biological function associated with labeling. However, SRS microscopy has limited specificity. This limitation is particularly relevant when imaging in the high wavenumber “CH” region (~2800 – 3500 cm^−1^). High wavenumber SRS imaging is desirable due to its high signal-to-noise ratio (SNR), but it is not as chemically specific as the “fingerprint region” of the Raman spectrum (~400-1700 cm^−1^). Nevertheless, the high-wave-number region still provides chemical information on lipids, proteins, nucleic acids, and water content in biological samples.^28–30^ Deep learning methods have been used to improve the specificity of high-wavenumber SRS imaging. Recent publications have shown the ability to create semantic segmentation maps of organelles within cells^31^, classify cancer^32^, and predict organelle fluorescence images^33^, all from label-free SRS images in the CH region.

Here, we show that hyperspectral CH SRS imaging can classify lymphocytes when combined with a powerful deep learning model. Specifically, we show that we can accurately distinguish B cells from T cells and differentiate T-cell subsets, with performance significantly improving on previous Raman spectros-copy-based classification. We compare Vision Transformer (ViT) with traditional deep learning classifiers and show that ViT-based classification offers superior balanced performance for the more challenging task of T-cell differentiation by using self-attention to model long-range relationships among spatial-chemical features, complementing convolutional models that primarily learn local hierarchical image patterns. These capabilities provide a general framework for future applications of label-free immune cell profiling in live tissue and immune engineering, where staining is undesirable or incompatible with live imaging.

## MATERIALS AND METHODS

### Lymphocytes Extraction and Separation

Male and female mice, 8-12 weeks of age, were used in compliance with University of Washington Institutional Animal Care and Use Committee Guidelines. We harvested spleens and generated a single-cell suspension by compressing the spleen between frosted glass slides. This cell suspension was filtered through 50 µm mesh into FACS buffer (HBSS, 0.5% BSA, 10 mM HEPES, pH 7.4). Cells were centrifuged at 350g for 4 minutes, resuspended in 1X RBC lysis solution at room temperature for 5 minutes, centrifuged, and resuspended in FC Block (33% 2.4G2 supernatant in FACS buffer) and incubated on ice for 30 minutes. Cells were stained with antibodies listed in the antibody table (Table S1) for 15-20 minutes in FC Block on ice. Cells were resuspended in FACS buffer and sorted with a BD FACSymphony™ S6 (BD Biosciences) as follows: Naïve CD8: CD8^+^CD62L^+^CD44^−^, Central Memory CD8: CD8^+^CD62L^+^CD44^+^, Effector Memory CD8: CD8^+^CD62L^−^ CD44^+^, Naïve CD4: CD4^+^CD62L^+^CD44^−^, Central Memory CD4: CD4^+^CD62L^+^CD44^+^, Effector Memory CD4: CD4^+^CD62L^−^CD44^+^, B Cell: CD19^+^B220^+^CD4^−^CD8^−^. We transferred sorted cells to poly-L-lysine-coated customized multiwell slides made from 3D-printed polylactic acid (PLA) wells attached to a coverslip and sealed with polydimethylsiloxane (PDMS). Cells were seeded at 37.2°C in a 5% CO2 incubator for 1 hour for better attachment, followed by fixation for 15 minutes at room temperature with 4% Paraformaldehyde (Electron Microscopy Sciences), rinsed with PBS, and coverslipped.

### *In vitro* T cell exhaustion model

Naïve T Cell was isolated from suspended splenocytes using EasySep™ Mouse Pan-Naïve T Cell Isolation Kit (StemCell™ Technologies), with starting population defined as CD3^+^ CD62L^+^CD44^−^. Tissue-culture plates were pre-coated with 1.2 μg anti-CD3 (Cytek) and 0.6 μg anti-CD28 (Cytek) in 300 μL PBS per well in a 24-well plate overnight at 4 °C, one day before harvesting cells from mice. On day 0, isolated naive T cell were seeded into basal medium consisting of RPMI-1640 (Ther-moFisher) supplemented with 10% heat-inactivated FBS (Gibco), 1% penicillin-streptomycin-glutamine (Gibco), 2% HEPES (Quality Biological), 1% sodium pyruvate (Gibco), 1% non-essential amino acids (Gibco), and 55 μM β-mercaptoethanol (Gibco) in each well. Cells were cultured in medium containing 100 U/mL IL-2 (Peprotech) and 10 ng/mL IFN-α (Bio-legend) to promote chronic stimulation-associated exhaustion and were maintained at 37.2 °C with 5% CO2. Subsequent restimulation steps were performed on fresh anti-CD3 coated well without anti-CD28 co-stimulation. Cells were replated in fresh cytokine-containing medium every two days and partially sampled for flow cytometry and imaging every four days. The experiment lasted 8 days. We monitored antigen and exhaustion-associated markers using CD44/CD62L and PD-1/Tim-3 expression to characterize cell state by Attune NxT (Invitrogen) flow cytometer. Details of antibodies used are listed in Table S1.

### Fluorescence and SRS Imaging

Fluorescence images of immunolabeled samples were acquired using a Nikon C2 laser scanning single-photon confocal microscope mounted on a Nikon TE2000 inverted microscope with excitation at 405, 488, 561, and 640 nm. SRS microscopy imaging was performed utilizing a homebuilt system on the same microscope.^34^ A single femtosecond laser oscillator (FLINT FL2, LIGHT CONVERSION) was used both as the Stokes beam (1030 nm) with a 75.25MHz (f0) repetition rate and to drive a homebuilt optical parametric oscillator (OPO) to produce a tunable pump beam set to 791 nm. Stokes beam was modulated at 18.82 MHz (f0/4) by an electro-optical modulator (EOM). Stimulated Raman loss was detected. Spectral focusing with grating stretchers for both beams was used to achieve narrowband SRS excitation at 16 cm^−1^ (measured with DMSO solvent).^35^ SRS images were acquired using a 40x, 1.25 NA objective (Olympus, UPlanSApo Silicone Oil Immersion) with a 1.4 NA condenser (Nikon). Both beams were set to a power of 40 mW at focus.

### Image Processing

Lymphocyte images were background-subtracted and corrected for field curvature. Cellpose-SAM^36^ was used to segment cells from average projected SRS images and fluorescence images. Image registrations were done by line ROI in ImageJ. Due to the non-adherent nature of suspension cells, certain cells may not be stabilized even on Poly-L-coated coverslips. In those cases, nearest-neighbor cell centroid matching was used to match cells in both modalities. Logarithm-transformed fluorescence intensity sum of each cell was used for cell identity annotation by a similar gating workflow as flow cytometry. To prepare cell images for model training and inference, we cropped each cell by centering the cell in the middle of a 60 × 60 or 100 × 100 pixel square for cell type and cell functional state classification, respectively. Pixels outside the centered cell are masked to 0. SRS spectra for individual cells were determined by averaging cellular pixel intensities at each Raman transition in the hyperspectral stack.

### Deep Learning Model Implementation

All classification models were implemented in PyTorch. Each spatial-spectral image cube was independently normalized to [0, 1] by min-max normalization across the whole cube. A Vision Transformer (ViT) divides an image into patches and uses self-attention to learn relationships among different image regions^37^ (Figure 2), whereas a convolutional neural network (CNN) extracts features using local convolutional filters. Although CNNs effectively identify local patterns, they generally integrate information from spatially separated regions progressively through multiple convolutional layers. In contrast, self-attention enables ViT to incorporate global information from earlier layers.^38^ We therefore selected ViT for SRS hyperspectral image classification (Figure 2) as the task may depend on spatially distributed cellular and biomolecular features rather than only localized features. Normalized image cubes were divided into non-overlapping spatial patches and embedded as a sequence of input tokens. Each hyperspectral image patch contains both local spatial information and SRS spectral profile and is represented as a spatial-chemical token. Multi-head self-attention is used in the transformer encoder to model relationships among spatial-chemical-aware tokens. A learnable classification token is used to aggregate patch-level information into a cell-level representation, followed by a fully connected classification head, in which its output dimension is adjusted according to the number of biological classes in each task. Corresponding model settings for each task are listed in Table S2. Datasets were split randomly into training/validation/testing in a 70/10/20 ratio. Data size in each class is summarized in Table 1. Hyperparameter optimization was performed using Optuna.^39^ Macro-F1 score was used as the optimization objective to account for class imbalance in the seven-class fine lymphocyte subtype model. The best hyperparameter set was then used to train the final model for evaluation on the held-out test set.

**Table 1.** Sample distribution for each classification task.

| **Task** | **Class** | **Train** | **Validation** | **Test** | **Total** |
|---|---|---|---|---|---|
|  | T | 17596 | 2520 | 5025 | 25141 |
|  | B | 4066 | 582 | 1162 | 5810 |
| **Lymphocyte lineage** | **Total** | **21662** | **3102** | **6187** | **30951** |
|  | B | 4066 | 582 | 1162 | 5810 |
|  | CD4 T | 8815 | 1263 | 2517 | 12959 |
|  | CD8 T | 8781 | 1257 | 2508 | 12546 |
| **Lymphocyte subset** | **Total** | **21662** | **3102** | **6187** | **30951** |
|  | B | 4066 | 582 | 1162 | 5810 |
|  | CD4 CM | 5269 | 754 | 1505 | 7528 |
|  | CD4 EM | 3159 | 453 | 902 | 4514 |
|  | CD4 N | 387 | 56 | 110 | 553 |
|  | CD8 CM | 1782 | 255 | 509 | 2546 |
|  | CD8 EM | 1166 | 168 | 333 | 1667 |
|  | CD8 N | 5833 | 834 | 1666 | 8333 |
| **Fine Lymphocyte subtype** | **Total** | **21662** | **3102** | **6187** | **30951** |
|  | Naïve | 408 | 116 | 60 | 504 |
|  | Activated | 397 | 113 | 58 | 568 |
|  | Ex-hausted | 472 | 135 | 68 | 675 |
| **T cell functional state** | **Total** | **1277** | **186** | **364** | **1827** |

**Figure 1.**
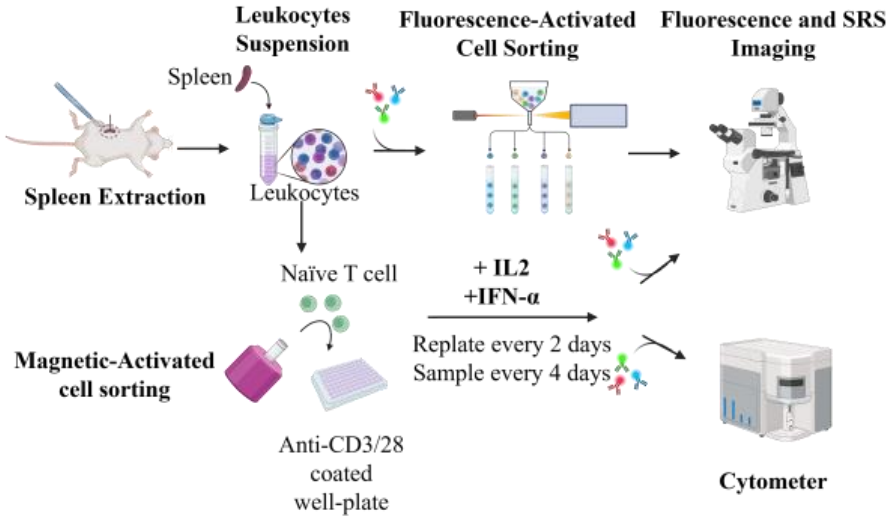
Schematic diagram of the experimental workflow. *Created in BioRender*.

**Figure 2.**
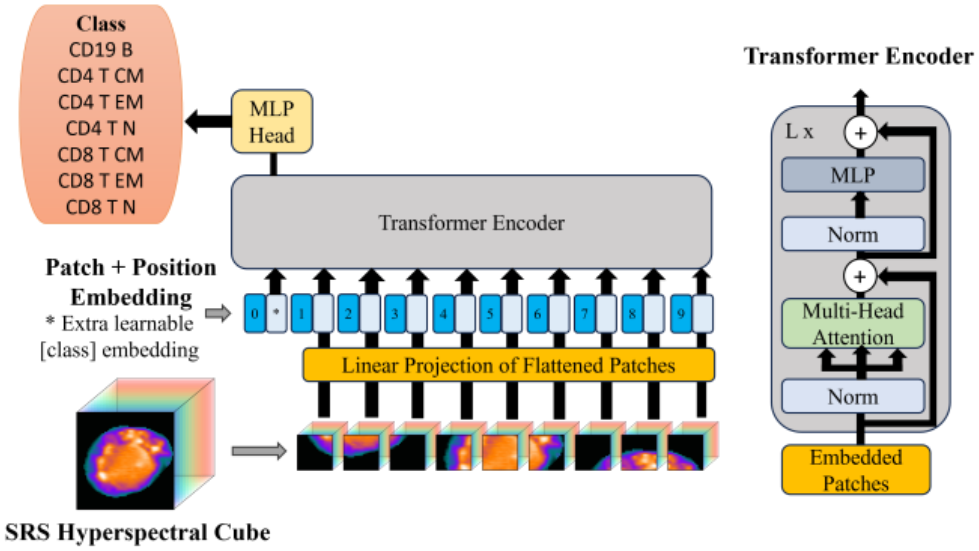
Schematic Diagram of Vision Transformer in SRS Hyperspectral Image Classification.

Conventional convolutional neural networks, such as ResNet-18, ResNet-50^40^, and VGG16 ^41^, were used as benchmark methods. All models were trained and evaluated using the same train/validation/test splits, input dimensions, preprocessing procedure, loss functions, class-balancing strategy, and evaluation metrics. The first convolution layer of CNN baselines was modified to accept full hyperspectral dimensions. Models were trained using the AdamW optimizer and cross-entropy loss. For imbalanced classification tasks, we adapted weighted random sampling to reduce bias toward highly represented classes. Online augmentation was applied to training only, including random combination of operations, such as random flip, rotation, spatial shifting, and additive Gaussian noise. Model performance was evaluated on held-out test data using accuracy, precision, recall, F1 score, receiver operating characteristic area under the curve (ROC-AUC), and confusion matrices. Models were trained with a Titan RTX GPU.

### Dimensionality Reduction and Representation Analysis

Uniform Manifold Approximation and Projection (UMAP)^42^ was used to visualize and compare the organization of different cell-level feature representations in order to assess how well conventional handcrafted features and model-learned representations captured cell differences. Three feature representations were compared, such as mean cellular Raman spectra, morphology features, and ViT extracted latent features. UMAP of Raman spectral features was performed using cosine distance to emphasize similarity in spectral profiles independent of overall intensity magnitude. Morphology features, including cell area, circularity, aspect ratio, solidity, eccentricity, major/minor axis length, and texture descriptors, were extracted from segmented and averaged projected SRS images and standardized before UMAP embedding in Euclidean space. For ViT, latent feature vectors were extracted from the final transformer layer before the classification head and embedded in Euclidean space.

## RESULTS

### Classification of Lymphocytes

We first demonstrate the capability of hyperspectral SRS microscopy in distinguishing B and T cells. The B and T cells were first separated via flow cytometry by marking CD4^+^ and CD8^+^ cells (T cells), CD4^−^ CD8^−^ CD19^+^ B220^+^ (B cells). The corresponding gating for cell identity is shown in Figure 3B. Representative SRS images of B and T cells in Figure 3A show no obvious distinguishing characteristics. Both T and B cells have large nuclei that occupy most of the cell area, with a high lipid signal in the cytoplasmic periphery, and similar cell size and roundness. When comparing normalized CH-SRS spectra (Figure 3C), CD4^+^, CD8^+,^ and B cells also highly overlap, with B cells showing a slightly stronger shoulder at 2880 cm^−1^ (asymmetric CH_2_ stretching). We further project normalized spectra into two-dimensional space using Principal Component Analysis (PCA), explaining 70% of the variance. However, B cells, CD4 T cells, and CD8 T cells all have strong overlap, as shown in Figure S1. By leveraging nonlinear manifold learning to preserve local neighborhood relationships while retaining aspects of broader data organization, UMAP produced relatively modest class separations, as shown in Figure S2. Nonetheless, similar to PCA results, CH spectra alone are insufficient to separate major lymphocyte types due to their broad features. Similarly, morphological feature-based UMAP cannot separate identities among them (Figure S3).

**Figure 3.**
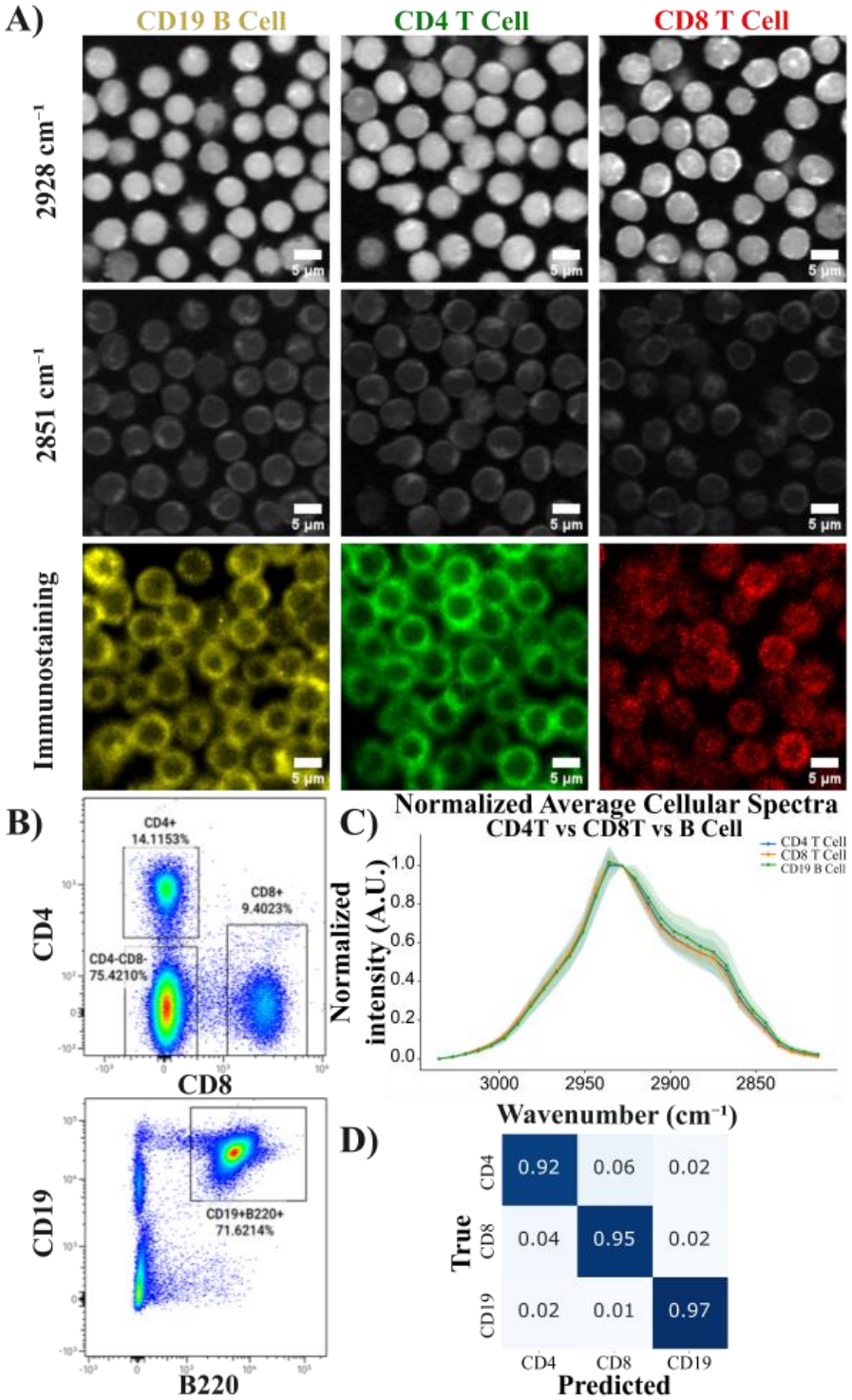
(A) Representative SRS images of CD4 T, CD8 T and CD19 B cell types with corresponding surface marker immunofluorescence imaging. Scale bar = 5 μm. (B) Representative Flow Cytometry Gating Scheme for separating B cells, CD4 T and CD8 T cells. (C) Average normalized cellular spectra for CD4^+^ T, CD8^+^ T and B cells. Standard deviations at each spectral point are depicted as the shading in each spectrum. (D) Confusion Matrix of Cell Type Classification in the broad Lymphocyte subsets testing set.

To leverage the spatial-chemical information for cell phenotyping, we used a powerful deep learning model, Vision Transformer, where a multi-head self-attention mechanism is applied to compare multi-channel intensity in each patch against other patches for learning spatially aware chemical patterns for each class. Figure 3D shows that ViT-based hyperspectral SRS classification performs well in separating CD4T/CD8T/B Cells, with overall classification accuracy at 93.8%, which shows higher accuracy than 60% in (CD4T/CD8T) classification in spontaneous Raman^24^ and 75.93% in (CD4T/CD8T/B Cells) classification in 3D refractive index tomography^43^. This result suggests that high spatial resolution images coupled with sub-cellular Raman spectral features help detect subtle differences between cell subtypes. Better classification performance using ViT compared to VGG-16 (92.3%), ResNet-18 (87.2%), and ResNet-50 (88.4%) motivates adoption of transformer-based architecture in further cell type classification tasks, as shown in Supplementary Figure 4.

### Classification of T Cell Subtypes

We next look at the more challenging task of classifying T cells in varying differentiation states, including naïve, effector memory, central memory CD4 T and CD8 T cells as shown in Figure 4A. After CD4 helper and CD8 cytotoxic T cell isolation by FACS, CD44 and CD62L markers further reveal their population within CD4 and CD8 T cells, as shown in Figure 4B. We identified naïve cells, CMs, and EMs based on markers (CD62L^+^CD44^−^), (CD62L^+^CD44^+^), and (CD62L^−^CD44^+^), respectively.

**Figure 4.**
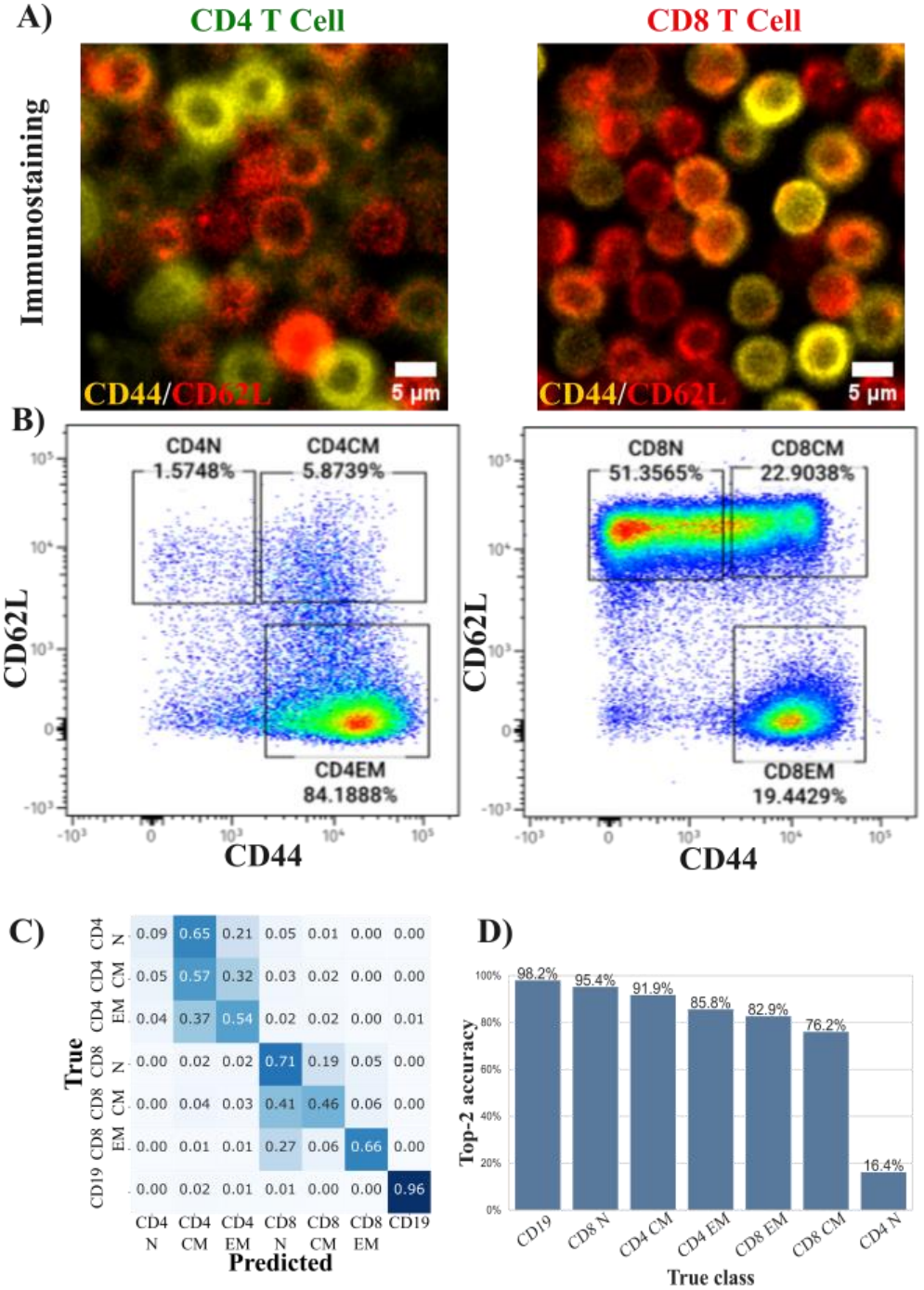
(A) Représentative CD44/CD62L immunofluorescence images of CD4 T, CD8 T cells. (B) Representative Flow Cytometry Plots of N/CM/EM T cell distribution in CD4 and CD8 T cells using the CD62L/CD44 antigen panel. (C) Confusion Matrix of Cell Type Classification in Fine Lymphocyte Subtypes testing set. (D) Corresponding Top-2 accuracy per class

The model performance is summarized in Figure 4C and Table S3. An overall accuracy (66%) and class-balanced accuracy (57%) are achieved, with all classes showing at least 46% accuracy except CD4 Naïve T cells. While more modest than the accuracy in distinguishing B and T cells above, we note this accuracy is notably higher than random guessing for the 7-class classification (14.3%). We attribute the low accuracy for classifying fine T cell subtypes in part to the limited CH spectral variation reflecting underlying biochemical differences. Moreover, the classification assumes a clear and true separation between classes; however, this assumption is not strictly accurate when based on thresholding of immunofluorescence intensity. Many cells could be defined as being in transitional states when near the border of the defined intensity gates, as seen in the flow cytometry plots (Figure 4B). As a result, the ground truth may be internally convolved, degrading accuracy in both model training and prediction. This hypothesis is supported by the analysis of the error. Most errors are caused by cells with neighboring surface marker expressions. When predictions are aggregated into the top-2 categories, the accuracy increases to 76 to 95% (Figure 4D), except for CD4N. We note that the poor CD4N result is likely due to its much lower representation in the dataset (Table 1), suggesting that augmented data acquisition may further improve model accuracy for the class.

### Classification of T Cell Functional State

Next, we imaged T cells under activation with hyperspectral C-H SRS to evaluate whether spatial-spectral features can distin-guish functional cell states. Morphological changes were clearly observed across the chronic stimulation window as shown in Figure 5A. At Day 4, average cell size increased 2-3-fold, with a higher cytoplasm-to-nucleus ratio compared to Day 0 isolated naïve T cells. Both Day 4 and Day 8 show increased lipid droplets in the intracellular regions, while Day 8 cells exhibit a relatively less homogeneous texture with more vesicle-like features compared to Day 4 cells. Flow cytometry analysis confirmed the progressive phenotypic transition of T cells during chronic stimulation as shown in Figure 5B. Isolated naïve cells (CD62L^+^CD44^−^) had no expression of PD-1 and Tim-3, as expected. Following stimulation by aCD3/CD28 and IFN-α^44^, the naive population shifted toward a CD62^−^CD44^+^ activated or effector-like phenotype with an increase in PD-1 expression^45^, while Tim-3 expression remained relatively limited on Day 4. Prolonged stimulation drove the cell population predominantly to CD62^−^CD44^+^ and PD-1^+^Tim-3^+^ double positive in Day 8. Co-expression of these inhibitory receptors indicates the development of an exhaustion-like phenotype^46^, which has been associated with more advanced T-cell dysfunction than PD-1 alone, including impaired proliferation and reduced production of IL-2, TNF-α, and IFN-’γ.^47^

**Figure 5.**
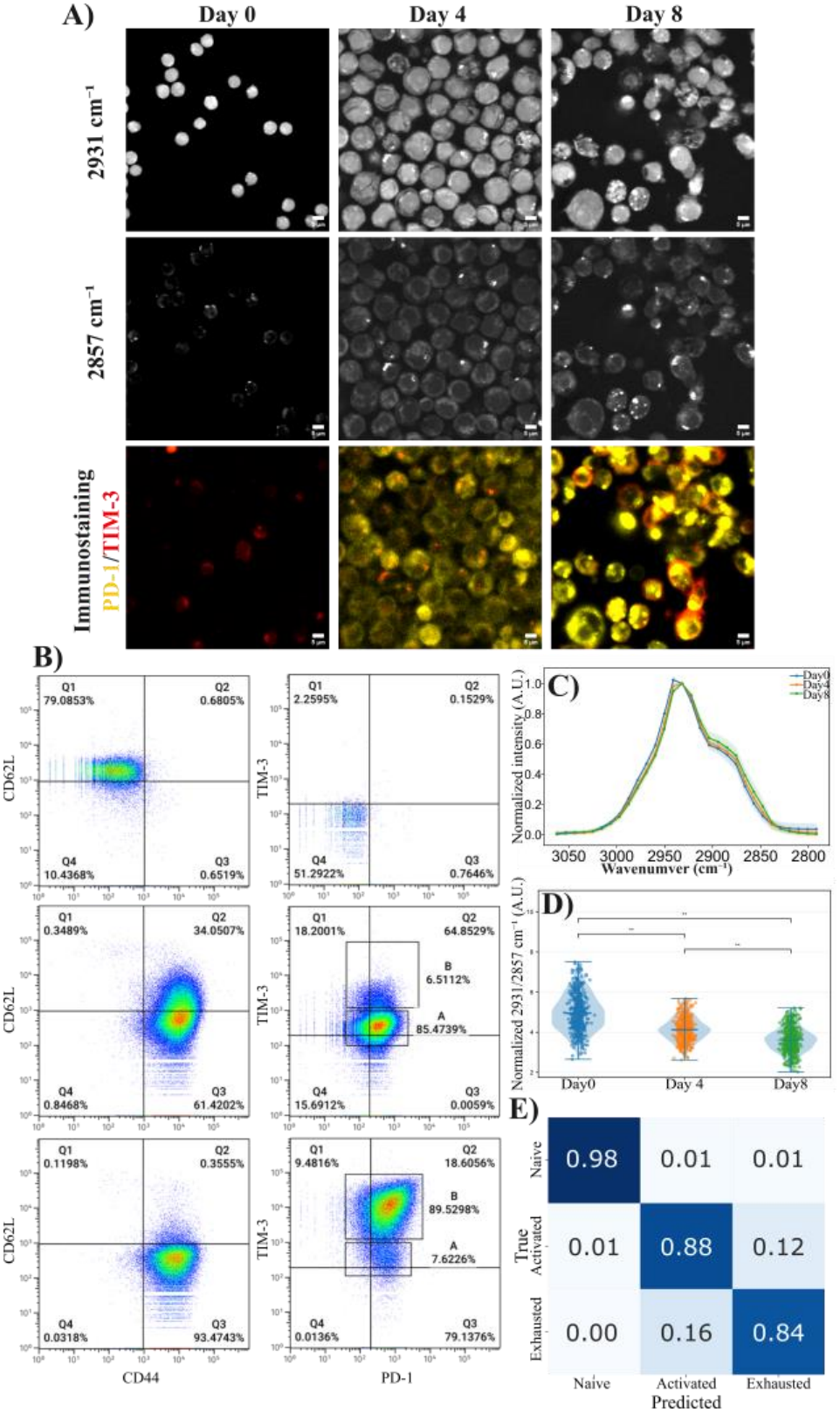
(A) Representative SRS and Tim-3/PD-1 immunostaining images of T cell functional state transitioning across Day0 to Day8. Scale bar = 5 μm. (B) Corresponding cytometry characterization in antigen (CD62L/CD44) and exhaustion panel (PD-1/Tim-3). (C) Average normalized cellular spectra for Day0, 4, and 8 (n = 500 each). Standard deviations at each spectral point are depicted as the area shading in each spectrum. (D) Normalized 2931/2857 cm^−1^ ratio across days. (E) Confusion matrix of Vision Transformer in naïve, activated, and exhausted-like T cell state classification.

Normalized average cellular spectrum (n=500 per day) shows subtle shape differences, with a slight shift towards lipids symmetric and asymmetric stretching as T cells progress from activation to exhaustion (Figure 5C). A progressive decreasing trend of the 2931:2857 cm^−1^ ratio shows statistically significant differences across days (p < 0.01), as shown in Figure 5D. The result is consistent with our observed increase in lipid droplet contents and previous literature reporting lipid remodeling that accompanies T cell activation.^48^ Activated T cells are also known to enlarge, proliferate, and generate new cellular membranes.^49^ One previous study has found that TCR/CD28 stimulation of CD4 T cells increased fatty-acid uptake, *de novo* lipid synthesis, and production of membrane lipids required for cellular enlargement and clonal expansion.^50^ Persistent antigenic stimulation induces mitochondrial stress and impaired oxidative phosphorylation^51^while dysfunctional CD8 T cells have been reported to accumulate cholesterol^52^, oxidized lipids^53^, and long-chain fatty acids^54^ that promote endoplasmic-reticulum stress, lipid peroxidation, and mitochondrial dysfunction.

To test our model’s ability to classify T cell functional states, we categorize Day 0, Day 4, and Day 8 cells as naïve T cells, activated T cells, and exhausted T cells, respectively. A separate ViT classification is trained using surface marker expression as the ground truth. The confusion matrix in the testing set shows accurate separation between naïve T cells and activated/exhausted cells, as shown in Figure 5E. However, there is around 12 to 16% cross-confusion between classifying activated and exhausted-like T cells, which is likely attributable to the 6 to 8 % of the cell population overlapping in the PD-1/Tim-3 panel, leading to ambiguous ground truth. The overall performance in T cell functional state classification reaches an accuracy of 89.84%, precision of 0.9, recall of 0.9011, F1 score of 0.9, and ROC-AUC of 0.9743.

Compared to fine-grained splenic T-cell subset classification, functional state classification shows much higher accuracy in the classification task. One possible explanation is that naïve, central-memory, and effector-memory T cells isolated from un-stimulated spleen are predominantly quiescent.^55^ Therefore, they potentially retain broadly similar bulk biochemical compositions. In contrast, activation and prolonged stimulation induce substantial metabolic remodeling^56,57^, generating stronger spatial-spectral contrast in the CH vibrational region.

### Latent Space Representation

We further examined the feature representations learned by ViT. UMAP was again used to embed ViT latent representation into 2D space. In comparison to the UMAPs based solely on spectral differences or extracted morphological features (Supplementary Figures 2 and 3, respectively), the embedded ViT feature space shows larger appreciable separation among many of the seven cell types, as shown in Figure 6A. Three main clusters were observed, which represent CD19 B cells, CD4 T cells, and CD8 T cells. Within the CD8 T cell cluster, three subtypes were partially separated, while only two subtypes were separable in the CD4 cluster. These results suggest that ViT transformed the original hyperspectral image cube information into a more class-discriminative feature space. The relative hierarchical confusion within classes (N/EM/CM within CD4 or CD8 clusters) is also well interpreted from the UMAP.

**Figure 6.**
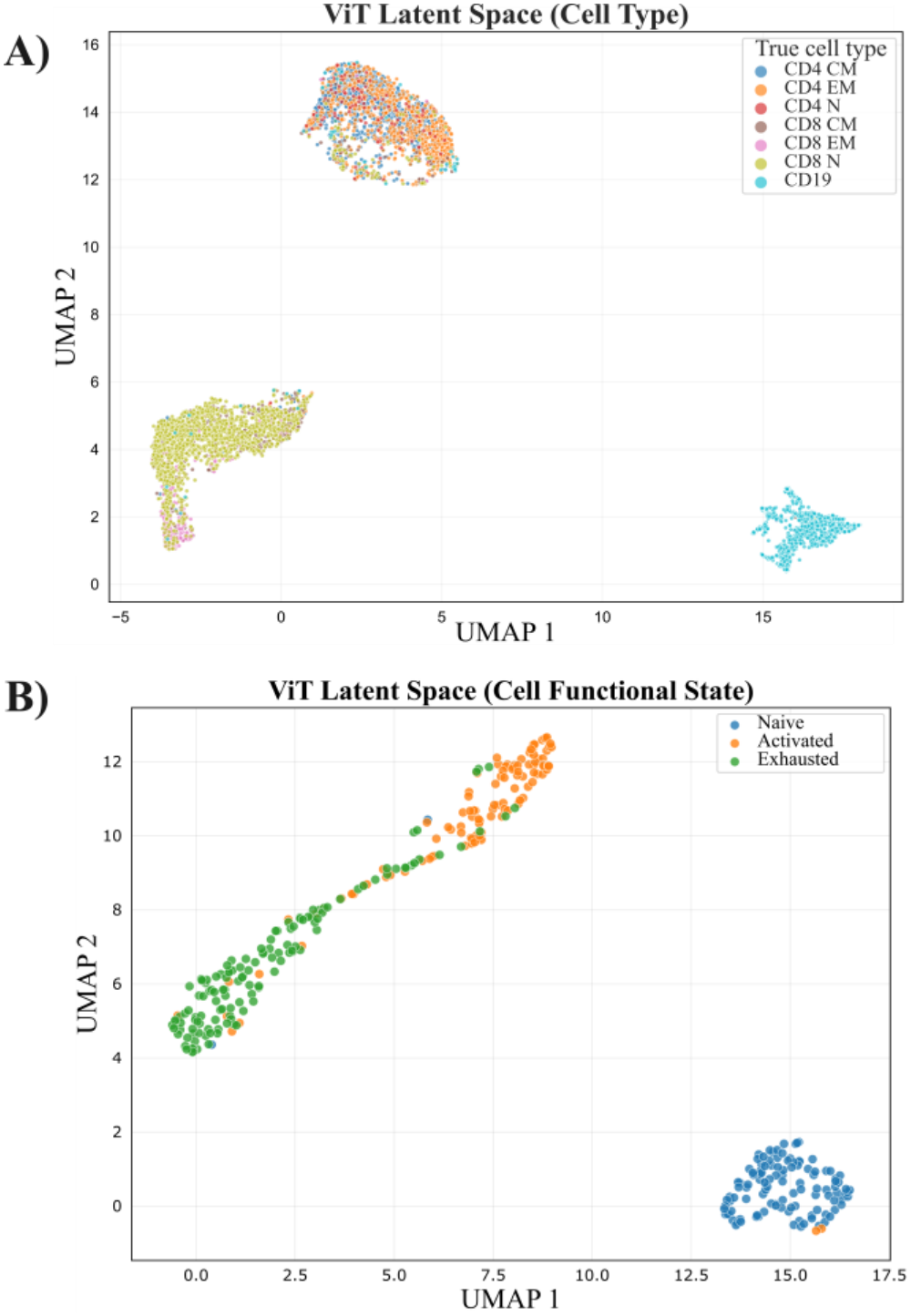
ViT Latent Feature Space Visualization in. (A) Fine lymphocyte subtypes (B) T cell functional states

Figure 6B shows a similar UMAP embedding in T cell function states. Here, ViT latent features show compelling separation between naïve, activated, and exhausted-like T cells. Interestingly, morphological features or spectral features alone also show these trajectory changes in UMAP with mostly separable classes (naïve T and activated or exhausted), though more modestly, as shown in Figure S5. Ultimately, this evidence suggests deep learning based on combined spatial-spectral features facilitates higher-accuracy modeling. Finally, we note that these clusters can potentially be used for monitoring state transitions as a continuum instead of discrete states. They can also be used to identify cell states based on cluster proximity, while out-of-distribution or abnormal cell types are marked for further characterization.

## DISCUSSION

Cell classification is essential for understanding cell function, differentiation, cell interaction, and drug response. T cells are an integral part of the adaptive immune system. The various types of T cells (e.g., cytotoxic, helper, regulatory) and their respective functions modulate dynamically throughout the life of a given cell and are an important point of study in immune response, vaccine development, and cancer treatment.^58^ Current methods for studying T cells typically rely on extraction, immunofluorescent antibody staining tagged to identify surface proteins, and separation via flow cytometry. While these methods can readily identify and collect the varying T cell differentiations, they are not applicable to *in situ* study of T cell function and response. Microscopy-based studies of T cells have recently shown promising results in understanding T cell selection and fate in situ, but they require fluorescent labeling, which may limit the observation breadth and duration.

In this study, we demonstrate an orthogonal label-free method for classifying immune cells from hyperspectral SRS images with ViT utilizing a self-attention mechanism. We successfully distinguish major lymphocyte populations, including CD4 T cells, CD8 T cells, and B cells, and further resolve naïve, central memory, and effector memory T-cell subtypes based on their intrinsic chemical signatures. Beyond subtype classification, we further investigate hyperspectral SRS imaging capability in characterizing dynamic cellular states associated with T-cell activation and exhaustion, demonstrating its potential to capture functional changes. These findings highlight the capability of hyperspectral SRS microscopy combined with deep learning to characterize both cell identity and functional states without exogenous labeling. This approach provides a complementary approach to conventional fluorescence-based profiling and establishes a foundation for investigating immune cell heterogeneity, activation, and exhaustion through label-free chemical imaging.

Although combining ViT and hyperspectral SRS imaging provides promising label-free multi-level immune cell type and state classification, model performance can be further improved by volumetric imaging to capture the full spatial-chemical organization rather than single z-plane imaging demonstrated here. We note that the single z-slice images here (~1 µm resolution) yield bias in signal and feature variation when imaging mixed cell types with different sizes in the same field of view. Longitudinal SRS imaging also provides temporal features and cellular-function descriptors to improve cell identity and state classification.^59^. We also note that biological states in single cells are potentially better described continuously rather than discretely.^60^

While immune cells are the focus of the study here, we expect that deep learning-augmented analysis of hyperspectral SRS images is a powerful tool towards classifying a wide variety of cell types and states at different levels. An exciting direction would be directly monitoring cell type distribution and motion in living tissue and animals. For cells with large morphological differences, SRS can already accurately identify them *in vivo*.^61^

We expect that the addition of chemical information would further enable finer classification of various cell types. In fact, it has recently been shown that Raman spectroscopy alone can predict the RNA profile of single cells.^62^ Future studies will focus on how much spectral and spatial information is needed to increase the discrimination of cell types and states in tissue. If necessary, either fingerprint SRS or Raman can be combined with morphological imaging to enable high-accuracy classification of cell types and cell states. This Raman phenotype-based approach complements other high-dimensional omics data approaches that can precisely define cell identity but lack the ability to track cells in 3D live tissue.

## ASSOCIATED CONTENT

### Supporting Information

Antibody specification; model training configuration; model performance comparison; PCA and UMAP embeddings using Raman or morphological features in cell type and function state; CNNs confusion matrix (PDF)

## AUTHOR INFORMATION

### Author Contributions

K.W.C. designed and performed the imaging experiments.

K.W.C. and P.L. designed and performed the extraction, isolation of splenocytes, culturing and chronic stimulation.

K.W.C. and Z.L. performed the deep learning analysis of the experimental results. B.M. performed initial pilot study and grant application. D.F. and H.Y.K. oversaw the project. The manuscript was written by K.W.C., B.M., and D.F. with contributions of all authors. All authors have given approval to the final version of the manuscript.

## ACKNOWLEDGMENT

This work was supported by funding from NIH R21AI183174 Award to D.F. The content is solely the responsibility of the authors and does not necessarily represent the official views of the National Institutes of Health.

