## Supplementary Information for "Label-Free Lymphocyte Subtype and State Classification by Stimulated Raman Scattering Microscopy"

**Content:**


- **Table S1:** Antibody Specification for Splenocyte Sorting or Cell State Characterization
- **Table S2:** Model Training Configuration in Fine Lymphocyte Subtype Classification
- **Table S3:** Deep Learning Model Performance Comparison in Fine Lymphocyte Classification
- **Figure S1:** Lymphocytes Cell Types Raman Feature PCA Embeddings
- **Figure S2:** Lymphocytes Cell Types Raman Feature UMAP Embeddings
- **Figure S3:** T and B Cell Morphological Feature UMAP Embeddings
- **Figure S4:** Confusion Matrices of Broad Lymphocytes Classification using CNNs Architectures
- **Figure S5:** T Cell Functional State UMAP Embeddings

| Marker | Fluorophore | Product Number | Manufacturer | Concentration |
|---|---|---|---|---|
| CD4 | BV510 | 100559 | BioLegend | 1:400 |
| CD8 | PerCp-Cy5.5 | 45-0081-82 | Invitrogen | 1:600 |
| CD19 | PE | 115508 | BioLegend | 1:600 |
| B220 | APC | 103212 | BioLegend | 1:1000 |
| CD44 | PE | 12-0441-82 | Invitrogen | 1:600 |
| CD62L | APC-e780 | 47-0621-82 | Invitrogen | 1:600 |
| PD-1 | PE | 12-9981-81 | Invitrogen | 1:100 |
| TIM-3 | APC | 119705 | BioLegend | 1:100 |

**Table S1:** Antibody Specification for Splenocyte Sorting or Cell State Characterization.

| Configuration | ViT | VGG 16 | ResNet-18 | ResNet-50 |
|---|---|---|---|---|
| Input size | 60 × 60 × 30 | 60 × 60 × 30 | 60 × 60 × 30 | 60 × 60 × 30 |
| Architecture | Vision Transformer | CNN | CNN | CNN |
| Patch size | 5 × 5 | N/A | N/A | N/A |
| Embedding dimension | 256 | N/A | N/A | N/A |
| Depth/ blocks | 8 Transformer blocks | 16 layers | 18 layers | 50 layers |
| Attention heads | 8 | N/A | N/A | N/A |
| Hidden/ MLP dimension | 2048 | 2048 / 1024 | 512 | 2048 |
| Dropout | 0.2424 | 0.2579 | 0.0877 | 0.3721 |
| Optimizer | AdamW | AdamW | AdamW | AdamW |
| Learning rate | $1.4294 \times 10^{-4}$ | $7.0722 \times 10^{-5}$ | $6.8162 \times 10^{-5}$ | $4.8895 \times 10^{-4}$ |
| Batch size | 128 | 64 | 64 | 128 |
| Epochs | 200 | 15 | 15 | 40 |

**Table S2:** Model Training Configuration in Fine Lymphocyte Subtype Classification.

| Model | Accuracy *(balanced)* | Macro Precision | Macro Recall | Macro F1 Score | ROC-AUC |
|---|---|---|---|---|---|
| ViT | 66.12% ***(56.81%)*** | 0.5597 | **0.5681** | **0.5622** | **0.8981** |
| ResNet-18 | 52.14% *(44.21%)* | 0.4715 | 0.4421 | 0.4323 | 0.8186 |
| ResNet-50 | 61.08% *(50.63%)* | 0.4889 | 0.5063 | 0.4865 | 0.8719 |
| VGG16 | **67.16%** *(47.52%)* | **0.5719** | 0.4752 | 0.4745 | 0.8975 |

**Table S3:** Deep Learning Model Performance Comparison in Fine Lymphocyte Classification.

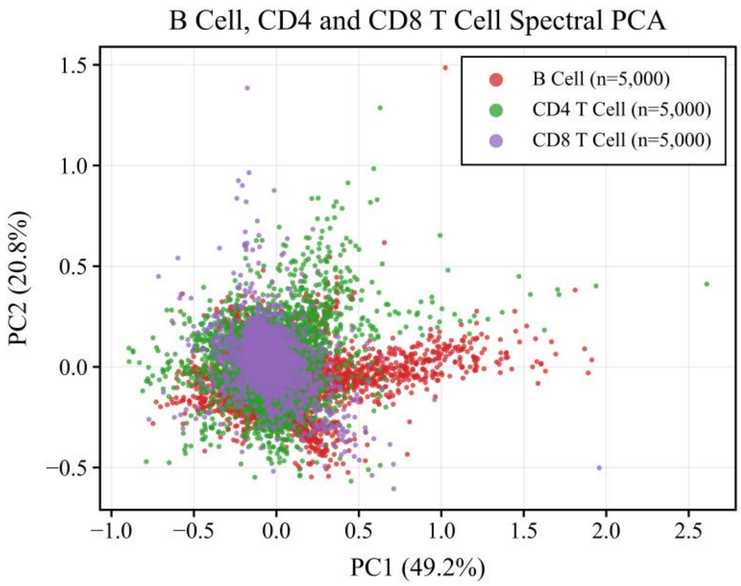


**Figure S1:** Lymphocytes Cell Types Raman Feature PCA Embeddings

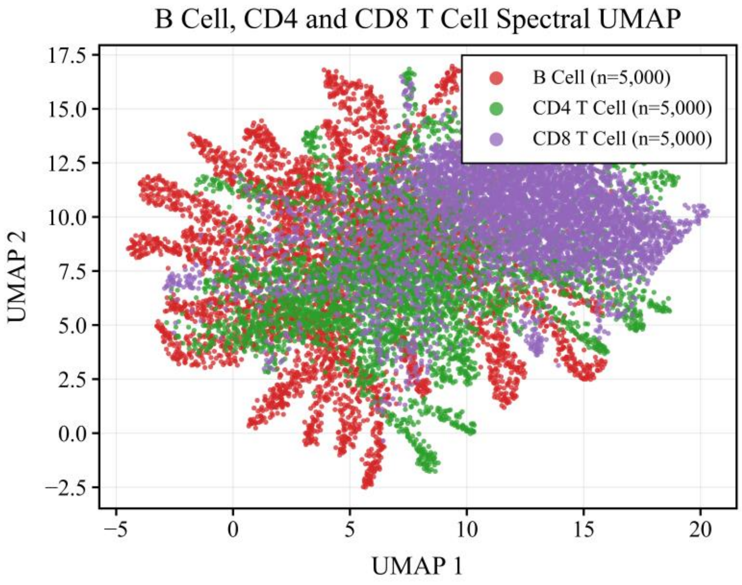


**Figure S2:** Lymphocytes Cell Types Raman Feature UMAP Embeddings

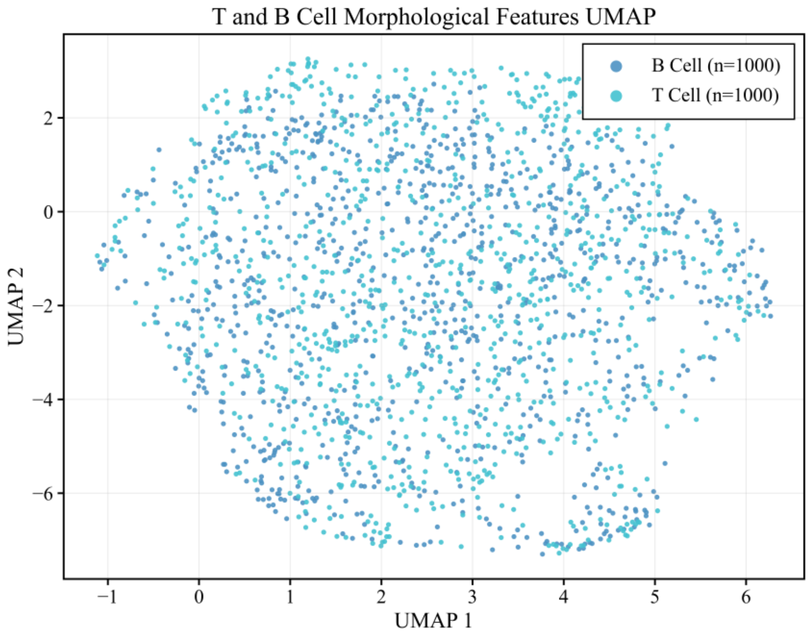


**Figure S3:** T and B Cell Morphological Feature UMAP Embeddings

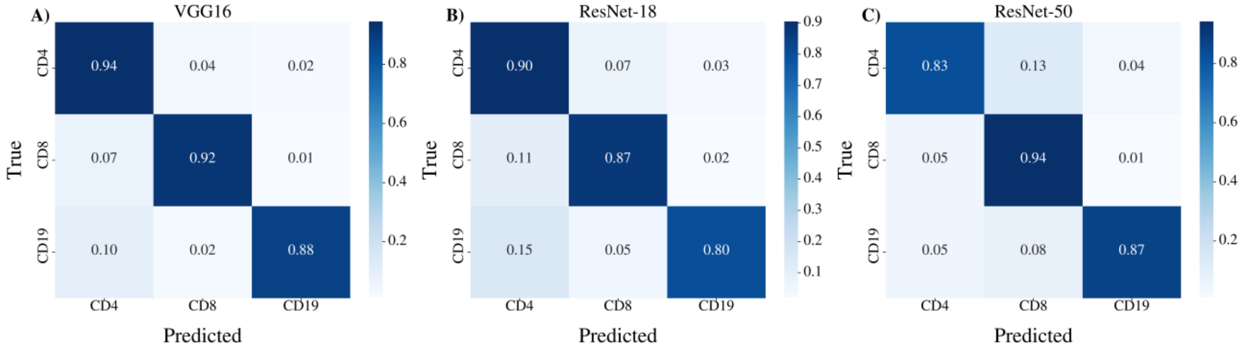


**Figure S4:** Confusion Matrices of Broad Lymphocytes Classification using CNNs Architectures. (A) VGG16. (B) ResNet-18. (C) ResNet-50.

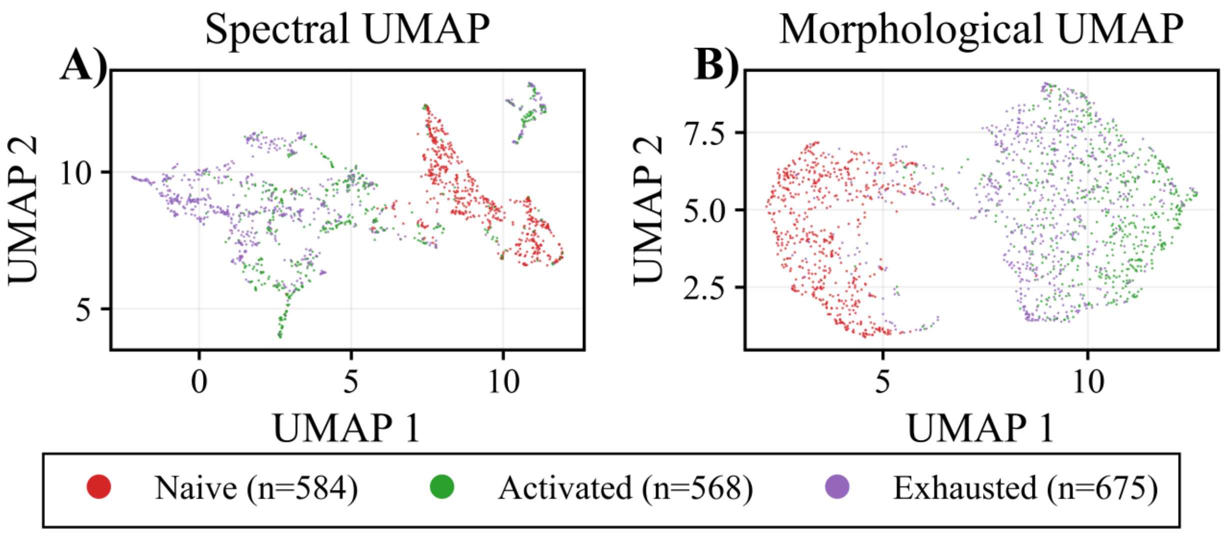


**Figure S5:** T Cell Functional State UMAP Embeddings. (A) using Morphological Features (B) Raman Features.
